# Behavioral flexibility, gut microbiome composition, and individual variability in oral oxycodone self-administration

**DOI:** 10.64898/2026.06.02.729613

**Authors:** Claire M. Corbett, Alexis E. O’Shall, Mark Niedringhaus, Elizabeth A. West

## Abstract

Prescription opioids such as oxycodone have been widely used in the United States and have contributed to the ongoing opioid epidemic. Although many individuals limit their use to prescribed contexts, a subset transitions to misuse and, in some cases, illicit opioid use. Identifying factors associated with variability in opioid intake may improve our understanding of vulnerability to opioid misuse. Here, we examined relationships among behavioral flexibility, gut microbiome composition, and subsequent oxycodone intake using an oral oxycodone self-administration model in male and female Long-Evans rats. Principal component analysis identified patterns of self-administration behavior that differentiated oxycodone-exposed animals from water controls. Among oxycodone-exposed animals, the principal component that contained variables of reinforced behaviors (i.e., active hole responding, reinforcers earned) was positively associated with oxycodone consumption, and hierarchical clustering identified three behavioral profiles distributed along this relationship. Behavioral flexibility, assessed before drug exposure using a contingency degradation task, was negatively associated with subsequent oxycodone intake, although it was not associated with the principal component measure of self-administration behavior. Oral oxycodone exposure was also associated with changes in gut microbiome composition, and analyses identified microbiome features associated with behavioral flexibility and oxycodone intake. Together, these findings demonstrate individual variability in oral oxycodone self-administration and identify behavioral and microbiome measures associated with subsequent drug-taking behavior. These results provide a foundation for future studies examining whether these measures prospectively contribute to vulnerability to opioid misuse.

## INTRODUCTION

Prescription opioids such as oxycodone are a main driver of the current opioid epidemic in the United States. While prescription opioids are a critical factor influencing the opioid epidemic, many individuals go no further than prescription use while a subset of individuals begin illicit use of prescription opioids and/or transition to heroin or fentanyl^1-5^. Notably, prescription opioids such as oxycodone are commonly administered and consumed orally in humans. However, only a small number of rodent preclinical studies have utilized operant oral oxycodone self-administration paradigms^6-8^ as the majority of preclinical oxycodone self-administration studies have utilized intravenous short (2 hours)^9^ or long (6 hours)^10-18^ access oxycodone self-administration paradigms. As such, there is little known about oral oxycodone self-administration, especially under an increased schedule of reinforcement. Additionally, as only a subset of individuals transition to illicit drug use, identifying potential associations between inherent behavioral and biological factors prior to drug use and future oral oxycodone taking behaviors in rodent models can provide insight into relevant preventative therapeutic measures prior to prescription. Prior work has shown that a history of drug use impairs goal-directed behavior in behavioral flexibility tasks^19-22^, however the relationship between intrinsic differences in behavioral flexibility^23^ and/or biological factors^24,25^ prior to drug use and future drug taking remains less characterized.

An emerging mediator of behavioral flexibility^26,27^ and substance use disorders^28-36^ is the gut microbiome. Dysbiosis of the gut microbiome due to prior drug use has been linked to substance use disorders in humans^37-40^ and preclinical studies using rodents^28-36^. Notably, these studies either investigated the effects of drugs of abuse^24,28-31^ on the gut microbiome or used antibiotics to decrease the gut microbiome in intravenous or experimenter-delivered drug studies prior to drug administration^32-36^. However, relatively little is known about how baseline gut microbiome composition, in the absence of antibiotic pretreatment, is associated with subsequent oral oxycodone self-administration or how the microbiome changes across the course of volitional oxycodone intake. Gut microbiome composition has been associated with performance on behavioral flexibility tasks, with microbiome dysbiosis via antibiotic delivery associated with poorer performance^26,27^. Accordingly, the present study examined whether baseline gut microbiome composition was associated with individual variability in behavioral flexibility and subsequent oral oxycodone consumption, as well as how the gut microbiome changed following volitional oral oxycodone self-administration. Next, we utilized the oral oxycodone self-administration paradigm to investigate the potential of a behavioral flexibility task^41^ as a behavioral biomarker of future oral oxycodone intake. Additionally, we investigated the overall effects of oral oxycodone on gut microbiome bacteria levels and the relationship among baseline gut microbiome composition, behavioral flexibility, and oral oxycodone intake.

## MATERIALS AND METHODS

### Subjects

Male (weight upon arrival: 225-300 g, aged 12-15 weeks old at start of self-administration) and female (weight upon arrival: 175-225 g, aged 12-15 weeks at start of self-administration) adult Long-Evans rats were purchased from Envigo (Indianapolis, IN, USA). Animals were maintained on a reverse 12:12 hour light/dark cycle (lights off at 9 A.M., lights on at 9:00 P.M.). Animals were singly housed and were provided with no enrichment throughout the duration of the experiment. Animals were randomly assigned to water or oxycodone groups. During nose poke training and initial oral oxycodone self-administration (self-administration days [SAD] 1-4), all animals were water regulated. All animals were maintained at no less than 85% of their free-water body weight by regulating water access in their home cage such that animals were not allowed to drop below 85% of their free-water weight as per standard fluid regulation guidelines (females: 17.5-22.25 mL of water; males: 20.5-25.5 mL of water; [*ad libitum* food]). Following completion of self-administration session 4, all animals were given free home cage access to water. We have previously reported contingency degradation data from a subset of rats^41^. We have not previously reported the oral oxycodone self-administration or gut microbiome data (water, n = 23, 8 males and 15 females; oral oxycodone, n = 36, 17 males and 19 females, 2 animals (1 male and 1 female) in the oxycodone group were not included in the contingency degradation due to equipment malfunction). We collected fecal matter from a subset of these animals (water, n=12, 6 males and 6 females; oral oxycodone, n=20, 10 males and 10 females). All procedures were approved by the Rowan University Institutional Animal Care and Use Committee (IACUC) and conducted in accordance with the US Public Health Service Guide for the Care and Use of Laboratory Animals.

### Behavioral training and testing

Animals were trained and tested in standard Med Associates operant chambers. All chambers were equipped with two spatially distinct levers with unilluminated cue lights above and an internal food receptacle between the spatially distinct levers on one side of the chamber that was used for all operant lever training for sucrose pellets, contingency degradation training, and contingency degradation test day. All chambers were also equipped with two spatially distinct nose poke holes with cue lights and an internal liquid trough located between the nose poke holes on the opposite side of the chamber, which was used for all operant oral oxycodone self-administration sessions. During the contingency degradation portion of the experiment, responses in the nose poke holes were without consequence. During the operant oral oxycodone self-administration portion of the experiment, levers remained retracted.

#### Contingency Degradation

Animals were first assessed for their behavioral flexibility using a contingency degradation task^41-45^ to determine if performance on this task is associated with future oral oxycodone intake. In this task, an action (i.e., left lever press) was degraded, and a different action (i.e., right lever press) remained intact after lever pressing was established. An animal’s ability to update this information in response to changes in their environment provides us with a measure of an animal’s inherent behavioral flexibility. First, animals underwent magazine training for 2 days to receive dustless precision sucrose pellets (Bioserve, Flemington NJ; males: 45 mg; females: 20 mg) into the internal food receptacle over 30 trials with a variable intertrial interval. Next, during operant lever training for sucrose pellets, animals underwent one training session per day (68 minutes), with each session consisting of four 15-minute blocks with a two-minute inter-block interval. During each block, one lever was extended. The lever extended was alternated between the two levers in each block (e.g., Block 1-left, Block 2-right, Block 3-left, Block 4-right, left and right as the starting lever, counterbalanced). Rats were first trained on a fixed ratio (FR) 1 schedule until they earned 50 pellets per lever over two days, followed by FR5 training (50 pellets per lever). Lastly, animals were trained on a variable ratio (VR) 9 schedule (range, 5-13 presses) for 4-5 days. Contingency degradation training consisted of a daily session (68 minutes) for 4 days. One of the two levers was randomly selected as the “contingent” or “non-degraded” lever and outcome delivery (sucrose pellet) was dependent upon the lever press under a VR9 schedule of reinforcement, while the other lever was selected as the “non-contingent” or “degraded” lever and outcome delivery did not depend on the animal’s actions (sucrose pellets were delivered every 30-40 seconds). On the contingency degradation test day, animals were tested under extinction for their behavioral flexibility (i.e., ability to learn and update the new information of lever contingencies). Both levers were extended, and lever presses were recorded but did not result in the delivery of a pellet for 15 minutes.

#### Oral oxycodone or water self-administration

Oral self-administration was conducted in Med Associates operant chambers that were distinct from those used for contingency degradation testing but had the same spatial orientation as described above. Depending on the fixed-ratio (FR) schedule, responses in the active hole resulted in delivery of water or oral oxycodone into the trough and, during self-administration sessions, illumination of a cue light within the active hole for 20 seconds. Responses in the inactive hole had no programmed consequences and were recorded as measures of response selectivity and nonspecific operant behavior. An automated syringe pump located outside each chamber (PHM-100VS-2, Med Associates) delivered liquid through tubing connected to the trough. Each earned reinforcer delivered 0.10 mL for males or 0.07 mL for females. Any liquid remaining in the trough at the end of a session was collected and measured. Any remaining volume was subtracted from the total volume delivered to calculate the volume consumed and, for oxycodone-treated animals, oxycodone intake in mg/kg.

First, animals were water regulated prior to nose poke training (Fig 1). Rats first completed three daily nose poke training sessions under an FR1 schedule, during which each active hole response resulted in the delivery of water without illumination of the cue light. Responses in the inactive hole had no programmed consequences. Following nose poke training, rats were split into water (n = 23, 8 males and 15 females) or oral oxycodone (n = 36, 17 males and 19 females) self-administration groups and underwent self-administration for 2 hours/day, 5 days/week, with two intervening days off, for a total of 21 self-administration sessions (Fig. 1). There was no difference in contingency degradation between the two self-administration groups (unpaired t-test, t(55)=1.543, p=0.129; water, n=23; oxycodone, n=34, 2 animals [1 male and 1 female] did not undergo contingency degradation due to equipment malfunction). During self-administration, earned reinforcers were accompanied by illumination of the active hole cue light and followed by a 20-s timeout period.

**Figure 1.**
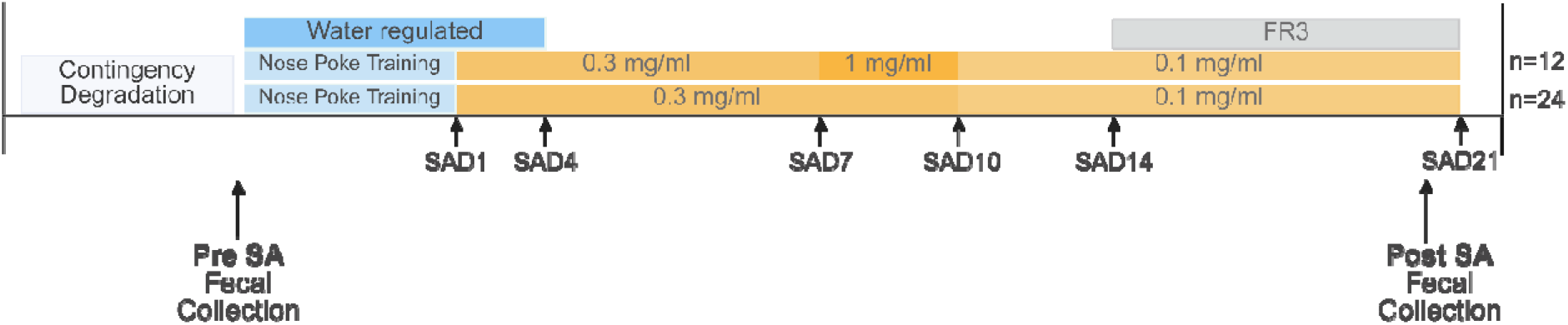
Experimental timeline. Animals underwent contingency degradation training prior to self-administration to assess behavioral flexibility. Prior to self-administration, all groups were water regulated prior to self-administration (dark blue) and underwent nose poke training (light blue) to obtain a water reinforcer prior to the start of oral oxycodone or water self-administration. All oral oxycodone animals self-administered oxycodone at a dose of 0.3 mg/ml from self-administration day (SAD) 1 to 6. A subset of animals (n=12) self-administered oral oxycodone at a dose of 1 mg/ml from SAD7 to 10 while most of the animals (n=24) self-administered oral oxycodone at a dose of 0.3 mg/ml from SAD7 to SAD10. All groups self-administered oral oxycodone at a dose of 0.1 mg/ml from SAD11 to SAD21. All groups were moved to a fixed-ratio (FR) 3 schedule of reinforcement on SAD14 to SAD21. Fecal samples were collected prior to self-administration (Pre) and during the last days of self-administration (Post).

During self-administration sessions 1-4, all animals remained water regulated and responded under an FR1 schedule. Oxycodone-treated animals self-administered 0.3 mg/mL oxycodone. Free home cage access to water was restored after completion of session 4. During sessions 5 and 6, all animals continued responding under the FR1 schedule, and oxycodone-treated animals continued to self-administer 0.3 mg/mL oxycodone^6^.

During self-administration sessions 7-10, animals remained on the FR1 schedule. A subset of oxycodone-treated animals (n = 12) self-administered 1.0 mg/mL oxycodone and were again regulated during these four sessions to promote responding for the reinforcer. The remaining oxycodone-treated animals (n=24) continued to self-administer 0.3 mg/mL oxycodone with free home cage access to water. The higher concentration was used to determine whether greater oxycodone exposure during early self-administration influenced subsequent responding for a lower concentration, however, we found no difference in the early exposure to oxycodone (Table S1), thus, combined the two groups for our analyses. Across sessions 1-10, sessions ended after 2 hours or after the animal earned 50-60 reinforcers, whichever occurred first, to limit the risk of excessive oxycodone intake during acquisition.

During self-administration sessions 11-13, animals continued responding under the FR1 schedule, but all oxycodone-treated animals self-administered 0.1 mg/mL oxycodone.^6^ Beginning with session 11, the reinforcer limit was removed, and animals were permitted to respond for the full 2-hour session regardless of the number of reinforcers earned.

During self-administration sessions 14-21, the response requirement was increased to FR3, such that three active hole responses were required to earn each reinforcer. All oxycodone-treated animals continued to self-administer 0.1 mg/mL oxycodone, and all animals were permitted to respond for the full 2-hour session.

### Estrous cycle analysis

We investigated the effects of the estrous cycle on the last 4 days of oral oxycodone self-administration. Additionally, since self-administration occurred 5 days/week with 2 days off, we did not capture proestrus in some of the females during the last 4 days of self-administration and they were excluded from this analysis (n=11). We examined the number of active hole responses and intake in a within-subjects design (n=23; 11 water, 12 oxycodone) such that the active hole responses and intake were analyzed when an individual female was in proestrus compared to when that same individual female was in non-proestrus stages. We analyzed the estrous cycle data in females in the proestrus stage compared to non-proestrus stages based on previous studies indicating that there is decreased opioid self-administration and intake during proestrus compared to all other stages^9,46,47^.

### Fecal collection

Fecal samples were collected from a subset of animals (water: n=12, 6 males and 6 females; oxycodone: n=20, 10 males and 10 females) at two time points: prior to water or oxycodone self-administration (Pre) and during the final self-administration days (SAD19-21; Post). To avoid interfering with self-administration behaviors and drug-associations with non-operant chambers, fecal samples were collected either from the animal’s home cage on cage change day within 1 hour of fresh bedding to ensure the fecal samples were collected on the same day as excretion, or if the animal had not excreted within the 1 hour from time of cage change to start of self-administration, fecal samples were collected from the operant chamber immediately following that day’s self-administration, which was wiped with 70% ethanol prior to behavioral testing. Samples were collected with sterile tools and put into sterile 1.5 mL tubes, immediately put on dry ice, and then stored at -80°C. Fecal collection was adapted from standard protocols^48^ because samples were obtained while animals were actively undergoing oral oxycodone self-administration, and oxycodone-associated constipation limited the feasibility of collecting freshly voided samples.

### 16S rRNA Sequencing, processing and analysis

Samples were shipped on dry ice to Omega Bioservices (Norcross, GA) for processing and analysis. All samples underwent and passed quality control analyses. 16S rRNA sequencing was performed using V3-V4 primers to detect bacteria and were analyzed using NextGen Sequencing software. All data were compiled on Illumina BaseSpace for further analysis. Raw 16S rRNA gene sequencing reads were processed using a standard bioinformatics pipeline to generate high-quality taxonomic profiles. Briefly, sequences were quality filtered, denoised, and merged, and amplicon sequence variants (ASVs) were inferred. Taxonomy was assigned using a curated reference database, and ASVs were aggregated to the species level to generate a count-based feature table for downstream analysis.

### Statistical analyses

To determine whether exposure to the higher oxycodone concentration during early self-administration affected subsequent oxycodone intake, intake was analyzed using a two-way mixed ANOVA with self-administration day as the within-subjects factor and early concentration group (1.0 mg/mL versus 0.3 mg/mL during sessions 7-10) as the between-subjects factor. Because no effect of early concentration group was detected, the two oxycodone groups were combined for subsequent behavioral analyses.

For longitudinal analyses, the self-administration protocol was divided into three phases: sessions 1-4, during initial acquisition and water regulation; sessions 5–13, during continued FR1 responding; and sessions 14-21, during FR3 responding. Sex effects on nose poke responding were initially assessed separately within each phase using four-way mixed ANOVAs with hole (active versus inactive) and self-administration day as within-subjects factors and sex (male versus female) and self-administration group (water versus oral oxycodone) as between-subjects factors. Reinforcers earned were analyzed using three-way mixed ANOVAs with self-administration day as the within-subjects factor and sex and self-administration group (water versus oral oxycodone) as between-subjects factors.

Within each self-administration phase, nose poke responding was analyzed using a three-way mixed ANOVA with hole (active versus inactive) and self-administration day as within-subjects factors and self-administration group (water versus oral oxycodone) as the between-subjects factor. Reinforcers earned were analyzed using a two-way mixed ANOVA with self-administration day as the within-subjects factor and self-administration group (water versus oral oxycodone) as the between-subjects factor. Significant main effects and interactions identified by ANOVA were followed by Bonferroni-corrected multiple comparisons.

Behavior during the final four self-administration sessions (sessions 18-21) was summarized by averaging each measure across sessions for each animal. Active and inactive hole responding during this period were analyzed using a two-way mixed ANOVA with hole (active versus inactive) as the within-subjects factor and self-administration group (water versus oral oxycodone) as the between-subjects factor. Within-session patterns of responding were examined by dividing each 2-hour session into four 30-minute bins. Nose poke responses were then analyzed using a three-way mixed ANOVA with hole (active versus inactive) and time bin as within-subjects factors and self-administration group (water versus oral oxycodone) as the between-subjects factor. Reinforcers earned were analyzed using a two-way mixed ANOVA with time bin as the within-subjects factor and self-administration group (water versus oral oxycodone) as the between-subjects factor. Estrous cycle effects were examined when the same female was in the proestrus stage compared to non-proestrus stages during the last 4 days of self-administration. Nose poke responses were analyzed using a three-way mixed ANOVA with hole (active vs inactive) and cycle (proestrus vs non-proestrus) as the within-subject factors and self-administration group (water vs oral oxycodone) as the between-subject factors.

Difference scores were calculated for active hole responses and reinforcers earned by subtracting the mean across the first four self-administration sessions from the mean across the final four sessions. Positive values therefore indicated an increase from early to late self-administration, whereas negative values indicated a decrease. Difference scores were compared between water-and oxycodone-exposed animals using Welch’s two-sample t-test.

#### Principal component and hierarchical cluster analyses

To characterize individual differences in self-administration behavior, principal component analysis (PCA) was performed in JASP (version 0.95.4). The PCA included four animal-level behavioral variables: mean (days 18-21) active hole responses during FR3, mean (days 18-21) inactive hole responses during FR3, mean (days 18-21) reinforcers earned during FR3, and the active responses early-to-late difference score. Before PCA, each variable was standardized as a z-score using its sample mean and standard deviation so that differences in measurement scale did not disproportionately influence the component solution. PC1 and PC2 accounted for 71% and 21% of the total variance, respectively. PC1 scores were compared between water-and oxycodone-exposed animals using Welch’s two-sample t-test.

Among oxycodone-exposed animals, behavioral profiles were examined using agglomerative hierarchical clustering based on PC1 scores and mean oxycodone consumption (mg/kg) across self-administration days 18-21. Both variables were standardized as z-scores using the “scale features” option in JASP. Clustering was performed using Euclidean distance and Ward’s D2 linkage. The number of clusters was selected according to the Bayesian information criterion (BIC), with a maximum of 10 clusters evaluated, resulting in a three-cluster solution. Cluster assignments were used to characterize empirically derived behavioral profiles and were visualized in the original, unscaled PC1-by-oxycodone-consumption space. A simple linear regression was used to assess the relationship between PC1 scores and mean oxycodone consumption across days 18-21 among oxycodone-exposed animals.

#### Contingency degradation analyses

A degradation index was calculated to normalize lever pressing during contingency degradation testing: (Contingent Lever Presses-Non-Contingent Lever Presses) / (Contingent Lever Presses + Non-Contingent Lever Presses)^41^. The index ranged from −1 to +1. A value of +1 indicated that all responses were directed toward the contingent lever, a value of 0 indicated equal responding on the contingent and non-contingent levers, and a value of −1 indicated that all responses were directed toward the non-contingent lever. Higher positive values indicated greater preferential responding on the lever that remained contingent and, consequently, greater sensitivity to the degraded action-outcome contingency. Values at or below zero indicated no preference for the contingent lever. Simple linear regression was used to assess the relationship between the degradation index and PC1 scores, and between the degradation index and oxycodone intake (days 18-21). Simple linear regression was used to determine the relationships separately by sex.

#### Microbiome analyses

Simple linear regression was used to evaluate associations between observed species richness and contingency degradation. Changes in observed species richness were analyzed using a two-way mixed ANOVA with fecal-sample time point (before versus after self-administration) as the within-subjects factor and self-administration group as the between-subjects factor. Bonferroni-corrected planned comparisons were used to evaluate the prespecified hypothesis that observed species richness would change following oral oxycodone self-administration. Genus differential abundance analysis was performed in R using DESeq2 (oxycodone relative to water control in the post-treatment fecal samples). Library size normalization was performed internally by DESeq2 using size factor estimation. For each species, log_2_ fold change (log_2_FC) values were calculated to quantify differences in abundance between oxycodone-exposed and water control group. Statistical significance was assessed using Wald tests and resulting p-values were adjusted for multiple comparisons using the Benjamini–Hochberg procedure to control the false discovery rate.

Finally, genus-level analyses were used to evaluate associations with behavioral flexibility and oxycodone exposure. Associations between baseline genus-level relative abundance and degradation-index scores were assessed using Spearman correlations. Oxycodone-associated changes in genus-level relative abundance were assessed using paired Wilcoxon signed-rank tests of within-subject log_2_ fold changes from pre-to post-self-administration samples. Genera meeting a nominal *p*<0.05 threshold in each analysis were subsequently mapped across both dimensions to identify taxa associated with both behavioral flexibility and oxycodone-associated change.

## RESULTS

### Rats self-administer oral oxycodone, and oxycodone-exposed rats respond more on a fixed-ratio 3 schedule than water control animals

No effect of sex was found on active or inactive hole responding during FR3 (Fig. S1A, B; Table S2), reinforcers earned (Fig. S1B, E, Table S3) or oxycodone consumption (Fig. S1E). Thus, males and females were combined within each self-administration group for subsequent FR3-focused analyses. We found no differences in self-administration behaviors (nose poke responding, reinforcers earned) between animals self-administering water and oral oxycodone during the early self-administration when rats were on an FR1 schedule (Table S4 and Table S5). When the response requirement increased to FR3, oxycodone-exposed animals exhibited greater overall nose poke responding compared to water control animals across self-administration days 14-21 as shown in Fig. 2A (main effect of self-administration group, F(1,57)=4.873, p=0.031). During this period, oxycodone-exposed animals made more active hole responses, although they did not earn more reinforcers compared to water control animals (Fig. 2B; F(1, 57)=2.93, p=0.093; Table S4). Nose poke responding also varied across self-administration days and was greater in the active hole compared to the inactive hole, although the trajectory of responding across days did not differ by self-administration group. Nose poke responding and reinforcers earned were further assessed to determine within-session patterns of responding across the last four days of self-administration (self-administration days 18-21) by dividing each 2-hour session into four 30-minute bins. Oxycodone-exposed animals demonstrated increased nose poke responding and reinforcers earned compared to water control animals across self-administration days 18-21, and both self-administration groups had increased active hole nose poke responding and earned more reinforcers during the first 30 minutes of the 2-hour sessions (Fig. S2, Table S6 and Table S7).

**Figure 2.**
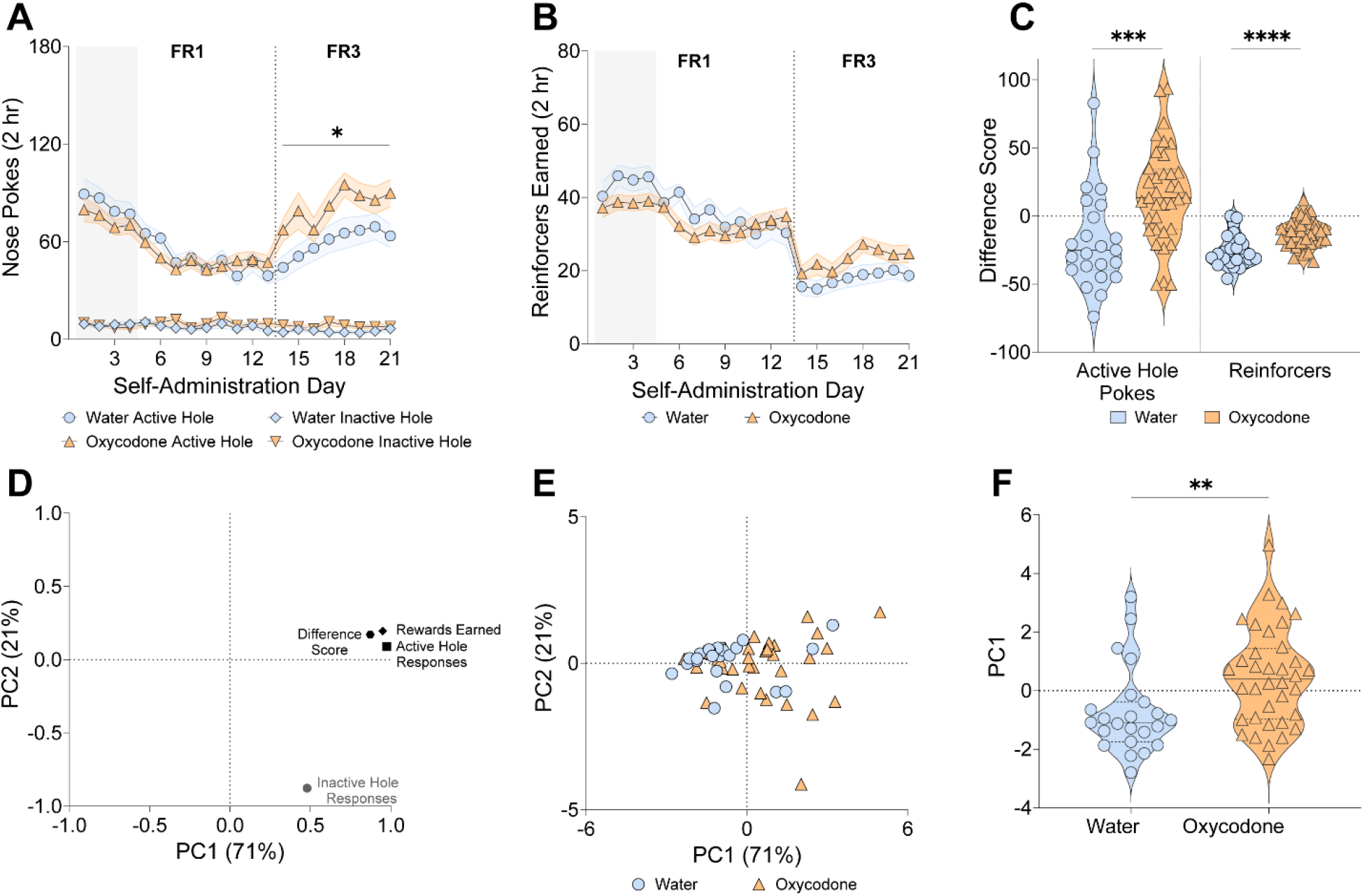
Oral oxycodone animals have increased active hole responding on a fixed-ratio 3 schedule compared to water controls. A. Nose pokes into the active and inactive holes of animals self-administering water (blue) or oral oxycodone (orange) during the 21 days of self-administration. Oral oxycodone animals have increased active and inactive hole nose pokes compared to water controls from self-administration day (SAD) 14-21 (three-way ANOVA; water vs oxycodone, F(1, 57) = 4.873, *p = 0.031), active hole vs inactive hole, F(1, 57) = 156.113, p < 0.001). B. Reinforcers earned during the 21 days of self-administration. C. Difference score of active hole nose pokes (left) and reinforcers earned (right) during the last four days of self-administration and first four days of self-administration. Oxycodone exposed animals have increased active hole (Welsch’s two-sample t-test, t(46.4)=3.46, *** p=0.001) and reinforcers earned (Welch’s two-sample t-test, t(41.7)=4.18, **** p<0.0001) difference scores compared to water controls. D. Principal component loading plot showing the contributions of active-hole responses, inactive-hole responses, reinforcers earned, and the early-to-late difference score to principal components 1 and 2 (PC1 and PC2), which accounted for 71% and 21% of the total variance, respectively. E. Individual PC1 and PC2 scores for water-exposed animals (blue circles) and oxycodone-exposed animals (orange triangles). Each symbol represents one animal. F. Oxycodone-exposed animals had higher PC1 scores than water-exposed animals (Welch’s two-sample t-test, t(50.96)=2.9, **p=.0057). Data are expressed as mean ± SEM. Water: n=23; oxycodone: n=36.

To quantify changes across self-administration, difference scores were calculated by subtracting each animal’s mean across days 1-4 from its mean across days 18-21. Oxycodone-exposed animals exhibited greater early-to-late increases in both active hole responding (Fig. 2C; Welch’s two-sample t-test, t(46.4)=3.46, p=0.001) and reinforcers earned compared to water control animals (Fig. 2C; Welch’s two-sample t-test, t(41.7)=4.18, p=0.0001). Thus, the treatment difference observed during late FR3 reflected a greater increase from initial responding among oxycodone-exposed animals rather than simply higher responding throughout self-administration.

The effect of the estrous cycle was assessed across self-administration days 18-21 where we found that oxycodone-exposed females consumed less oral oxycodone during proestrus compared to non-proestrus stages (Fig. S3C; paired t-test; t(11)=2.423, p=0.0338), however there was no effect of estrous cycle on nose poke responding across self-administration days 18-21 (Fig. S3A,B; Table S9).

To integrate multiple dimensions of self-administration behavior, we performed a principal component analysis using active hole responses, inactive hole responses, reinforcers earned, and the early-to-late difference score. PC1 accounted for 71% of the total variance and was characterized by strong positive loadings from active hole responding, reinforcers earned, and the difference score (Fig. 2D). PC2 accounted for an additional 21% of the variance and was distinguished primarily by the strong negative loading of inactive hole responding. Visualization of individual PC1 and PC2 scores revealed that oxycodone-exposed animals were generally shifted toward higher PC1 scores, whereas water-and oxycodone-exposed animals showed substantial overlap along PC2 (Fig. 2E). Consistent with this pattern, oxycodone-exposed animals had significantly higher PC1 scores than water controls (Fig. 2F; Welch’s two-sample t-test, t(50.96)=2.9, p=0.0057, Fig. 2F). Thus, oxycodone exposure was associated with a shift along a continuous behavioral dimension reflecting greater reinforced responding and a larger increase in responding across self-administration.

To further characterize individual differences among oxycodone-exposed animals, agglomerative hierarchical clustering was performed using PC1 scores and mean oxycodone consumption across self-administration days 18-21. Optimization according to the Bayesian information criterion supported a three-cluster solution. The solution accounted for 80.7% of the variance in the two clustering features (R^2^=0.81) and yielded a silhouette coefficient of 0.530, indicating moderate separation among the empirically derived behavioral profiles. The clusters were generally distributed along a continuum of increasing oxycodone consumption and PC1 scores (Fig. 3A). Cluster 1 was characterized predominantly by lower oxycodone consumption and lower PC1 scores, Cluster 2 occupied an intermediate region across the two measures, and Cluster 3 was characterized by higher consumption and higher PC1 scores. Mean oxycodone consumption was strongly and positively associated with PC1 scores across oxycodone-exposed animals (simple linear regression, R^2^=0.69, p<.0001). Thus, animals that consumed more oxycodone during late FR3 also generally exhibited higher scores on the behavioral dimension reflecting reinforced responding and early-to-late increases in responding.

**Figure 3.**
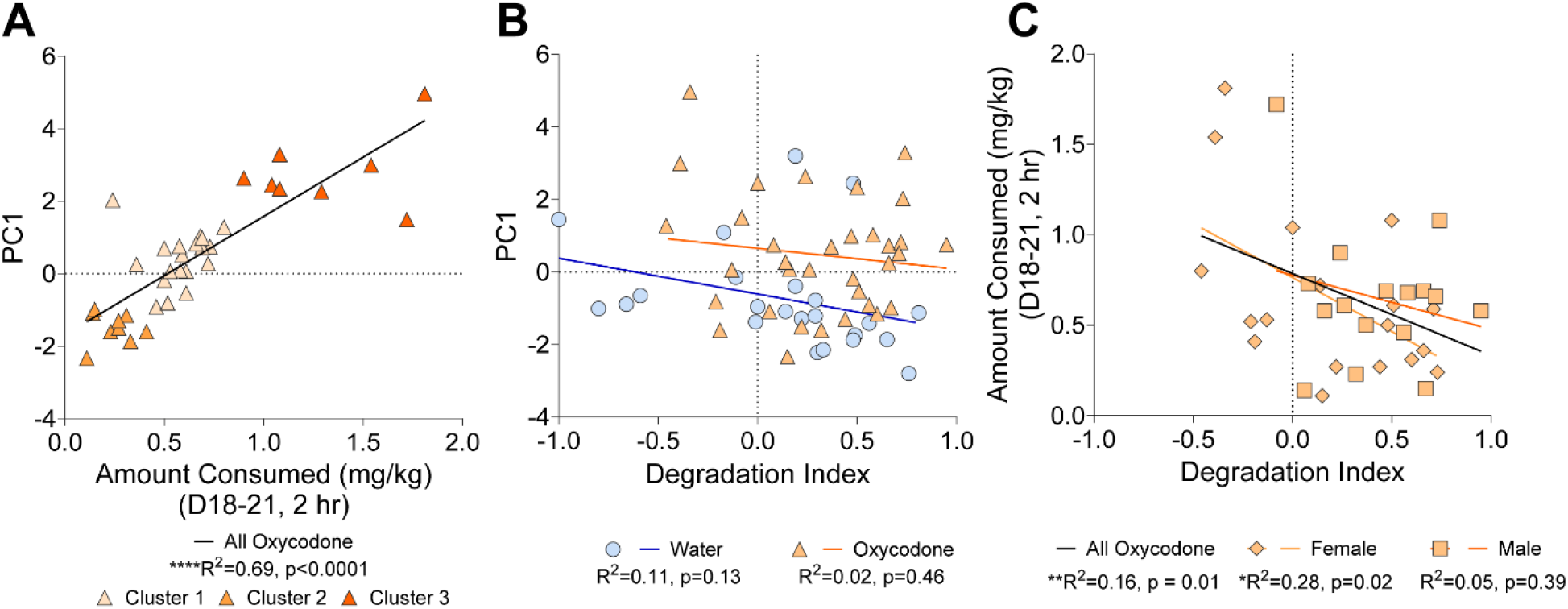
Oral oxycodone consumption is associated with principal-component scores and contingency-degradation performance. A. The amount consumed (mg/kg) of oxycodone is positively correlated with the principal component 1 score (PC1; simple linear regression, R^2^=0.69, ****p<0.0001) and hierarchical clustering identified three groups of oxycodone-exposed animals (n=19, 9, and 8; model: R^2^=0.807; BIC = 35.03; silhouette coefficient=0.530). B. Principal component 1 score is not correlated with performance on the contingency degradation task in water (blue) or oxycodone (orange) animals. C. The amount consumed (mg/kg) of oxycodone is negatively correlated with performance on a behavioral flexibility task as measured using a degradation index (simple linear regression, **R^2^ = 0.16, p = 0.01). The amount of oral oxycodone consumed by females is negatively correlated with performance on the contingency degradation task (*R^2^=0.28, p=0.02) but not in males (R^2^ = 0.05, p = 0.39). Water: n=23; Oral oxycodone: n=34; males: n=16, females: n=18.

### Performance on the contingency degradation task before drug exposure was associated with subsequent oral oxycodone consumption

Performance on the contingency degradation task was summarized using a degradation index calculated from contingent and non-contingent lever presses^41^. We first examined whether contingency degradation performance was associated with PC1, the broader dimension of self-administration behavior identified by PCA. Degradation index was not significantly associated with PC1 scores in either water-exposed animals (R^2^=0.11, p=0.13) or oxycodone-exposed animals (R^2^=0.02, p=0.46) as shown in Fig. 3B. We next examined whether contingency degradation performance was specifically associated with subsequent oxycodone consumption. Degradation indices were negatively associated with mean oxycodone consumption during self-administration days 18-21 (R^2^=0.16, p=0.01, Fig. 3C). Animals with higher degradation indices, indicating greater sensitivity to the degraded contingency, subsequently consumed less oxycodone during late FR3 self-administration. In sex-stratified analyses, the negative association was significant among females (R^2^=0.28, p=0.02) but not among males (R^2^=0.05, p=0.39). Thus, contingency degradation performance was associated with subsequent oxycodone consumption but did not explain variation in the broader multidimensional self-administration profile captured by PC1.

### Oral oxycodone self-administration disrupts the gut microbiome compared to water self-administration

To examine relationship between behavioral flexibility and the gut microbiome we first examined the number of species after the contingency degradation task and prior to self-administration (Pre) and found that the number of species negatively correlated with performance on the contingency degradation task (Fig. 4A; simple linear regression; F(1, 28)=9.7, R^2^=0.26, p=0.004). Our prespecified hypothesis that oral oxycodone self-administration would change gut microbiome species richness compared to animals self-administering water was tested by examining the number of species in the gut microbiome prior to (Pre) and at the end of (Post) self-administration in both water and oral oxycodone-exposed animals. We found a significant main effect of self-administration group (two-way ANOVA, (F(1, 30)=13.03, p=0.001) and no effect of time point (pre vs post, F(1, 30)=1.43, p=0.24) or self-administration group and time point interaction (F(1, 30)=3.20, p=0.08). Planned comparisons of self-administration group and time point showed a significant difference between the number of species prior to oral oxycodone exposure and post oral oxycodone exposure (p=0.042), where there was a decrease in the number of species after oral oxycodone exposure (Fig. 4B). Additionally, there was a significant difference between the number of species post self-administration between oxycodone-exposed and water control animals (p=0.008). Lastly, oxycodone exposure resulted in broad alterations in the gut microbiome at the species level. Differential abundance analysis revealed a greater number of species with significantly decreased abundance in the oxycodone group relative to water controls than the number of species that were increased (post-treatment), suggesting a general reduction in specific microbial taxa following oxycodone exposure (Fig. 4C).

**Figure 4.**
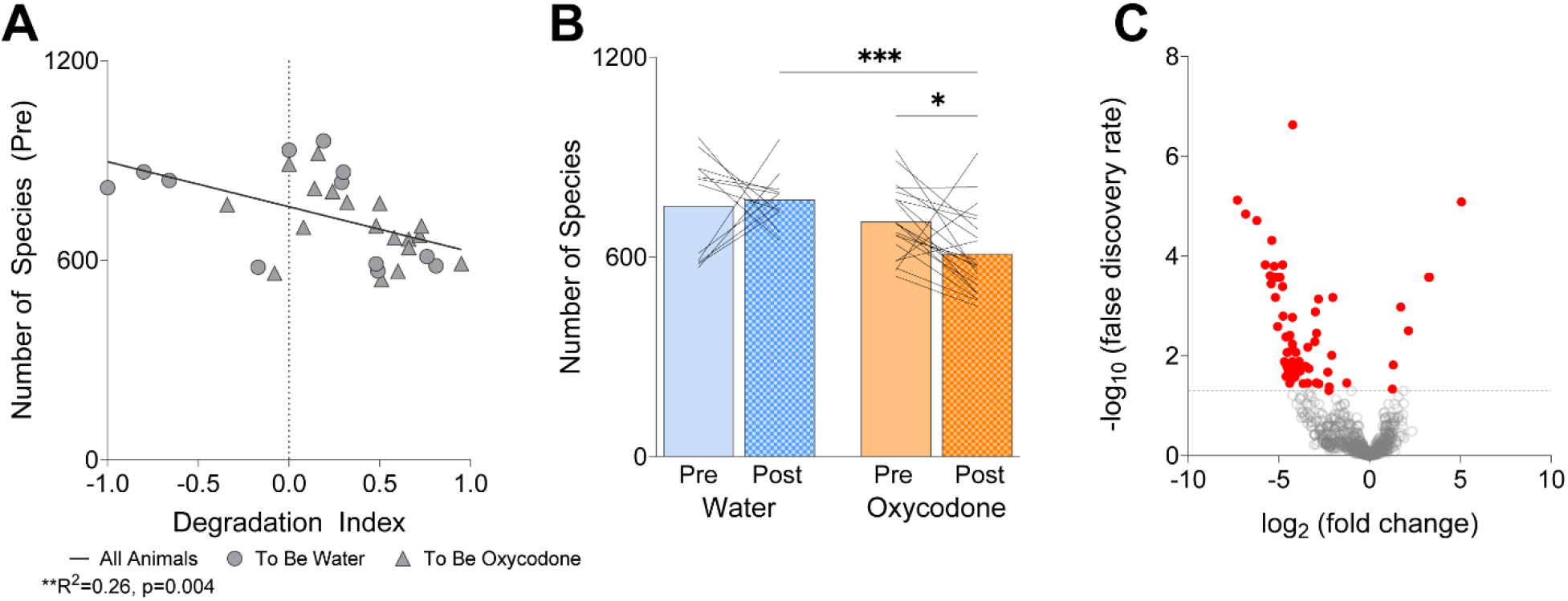
Oral oxycodone exposure decreases the number of species in the gut and correlates with the number of species in the gut following oral oxycodone exposure. **A.** Degradation index negatively correlated with the number of species in the gut following contingency degradation (prior to self-administration; simple linear regression; R^2^=0.26, p=0.004). **B.** There was a decrease in the number of species in the gut following oral oxycodone exposure (Post) compared to levels before drug exposure (Pre; planned comparisons, *p=0.042) and compared to the number of species following water self-administration (Post; planned comparisons, ***p=0.008) **C.** Volcano plot showing differential abundance of gut microbiome species following oxycodone exposure relative to water control. Red circles indicate species that are significantly different between water-and oxycodone-exposed groups (false discovery rate <0.05). Most significant species exhibit negative log_₂_(fold change) values, indicating decreased abundance following oxycodone exposure. Water: n=12; Oral oxycodone: n=20.

Finally, to determine whether specific microbial features were associated with behavioral flexibility and oxycodone exposure, we performed genus-level analyses examining relationships between relative abundance, behavioral flexibility, and oxycodone associated changes. Baseline relative abundances of 66 genera were nominally associated with behavioral flexibility (*p* < 0.05), with 14 genera showing positive and 52 showing negative Spearman correlations (Fig. 5A). Similarly, 70 genera showed nominal changes in abundance following oxycodone exposure (*p* < 0.05), with 10 genera increasing and 60 decreasing in abundance (Fig. 5B). To integrate these findings, genera were mapped according to both their oxycodone-associated change and their association with behavioral flexibility (Fig. 5C). Nine genera met the nominal *p* < 0.05 criterion in both analyses (Table 1). All nine genera decreased following oxycodone exposure. Baseline abundance of seven of these genera was negatively associated with behavioral flexibility, whereas baseline abundance of the remaining two was positively associated with flexibility.

**Figure 5.**
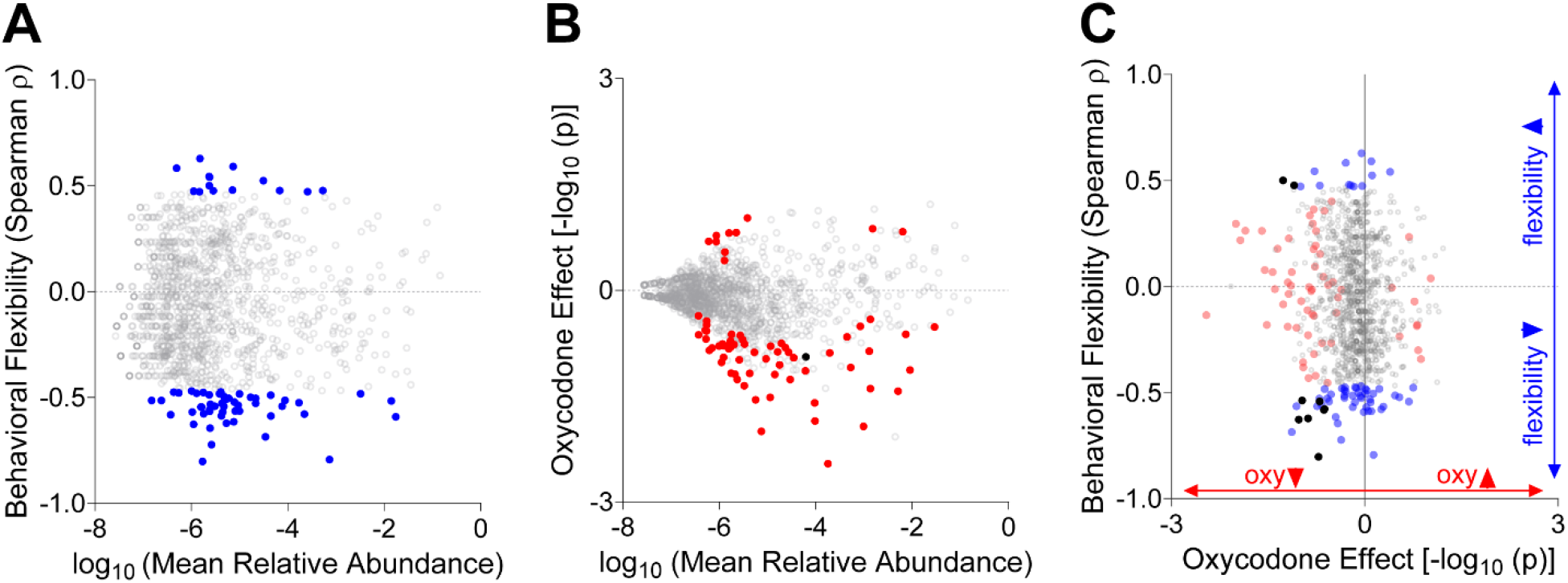
Genus-level relationships between relative abundance, oxycodone exposure, and behavioral flexibility. A. Relationship between genus-level mean relative abundance and behavioral flexibility (Spearman r). Genera showing nominal associations with behavioral flexibility (*p*<0.05) are indicated by blue circles (14 positive and 52 negative associations). B. Genus-level changes in relative abundance from pre-to post-self-administration in oxycodone-exposed animals. Genera showing nominal changes following oxycodone exposure (*p*<0.05) are indicated by red circles (10 increased and 60 decreased). Oxycodone effects are expressed as directional −log_10_ (*p*) values, with negative values indicating decreased and positive values indicating increased abundance following oxycodone exposure. Genera showing nominal changes following both oxycodone and water self-administration are indicated as black circles. C. Integrated analysis of genus-level oxycodone-associated changes and behavioral flexibility. Black circles indicate nine genera meeting the nominal *p* < 0.05 criterion in both the oxycodone and behavioral-flexibility analyses. All nine genera decreased following oxycodone exposure; baseline abundance of seven was negatively associated with behavioral flexibility, whereas baseline abundance of two was positively associated with flexibility. Water: n=12; Oral oxycodone: n=20.

**Table 1.** Genus taxa that are oxycodone sensitive and correlated with behavioral flexibility.

| **Genus** | **log<sub>2</sub> Fold Change Oxycodone** | **Spearman r** | **Relative Abundance** | **Interpretation** |
|---|---|---|---|---|
| *Oxalophagus* | -1.27E+00 | 5.00E-01 | 2.36E-06 | ↓ oxy / ↑ flexibility |
| *Jonquetella* | -1.10E+00 | 4.77E-01 | 5.35E-04 | ↓ oxy / ↑ flexibility |
| *Geothrix* | -1.02E+00 | -6.27E-01 | 1.12E-06 | ↓ oxy / ↓ flexibility |
| *Rhodopila* | -9.72E-01 | -5.37E-01 | 9.36E-06 | ↓ oxy / ↓ flexibility |
| *Phenylobacterium* | -8.79E-01 | -6.21E-01 | 5.35E-06 | ↓ oxy / ↓ flexibility |
| *Pontibacillus* | -7.18E-01 | -8.02E-01 | 1.72E-06 | ↓ oxy / ↓ flexibility |
| *Tannerella* | -7.00E-01 | -5.41E-01 | 3.05E-06 | ↓ oxy / ↓ flexibility |
| *Fontibacter* | -6.32E-01 | -5.81E-01 | 3.73E-07 | ↓ oxy / ↓ flexibility |
| *Actinomycetospora* | -6.26E-01 | -5.77E-01 | 1.80E-06 | ↓ oxy / ↓ flexibility |

## DISCUSSION

Many individuals are prescribed oxycodone and the majority of these individuals will terminate use upon completion of the prescription^49^. However, a subset of individuals prescribed oxycodone will misuse the drug^50,51^, indicating individual differences in the patterns of oxycodone use. Extended-access intravenous studies have demonstrated the emergence of distinct phenotypes during oxycodone self-administration^52-54^, however, individual variability during short-access oral oxycodone self-administration has been less extensively characterized. Here, we sought to characterize short-access oral oxycodone self-administration in both male and female rats compared to a water control group and found that water-and oxycodone-exposed animals had similar levels of responding for reinforcers under the FR1 schedule of reinforcement. Importantly, oxycodone-exposed animals increased responses into the active hole when the schedule of reinforcement was moved to FR3. Within-session responding during the last four days of self-administration demonstrated that oxycodone-exposed animals responded more and earned more reinforcers than water control animals and that both self-administration groups had increased responding and reinforcers earned during the first 30-minute interval of the 2-hour sessions. Oxycodone-exposed animals had higher principal component 1 (PC1) scores, which represented reinforced behavior, than water control animals. Among oxycodone-exposed animals, PC1 scores were positively associated with the amount of oxycodone consumed, indicating that greater expression of the self-administration behaviors represented by PC1 was associated with greater oxycodone intake. Hierarchical clustering based on PC1 scores and oxycodone consumption further identified three behavioral profiles distributed along this relationship. Identifying this variability in an oral oxycodone model is relevant as prescription oxycodone is typically ingested orally, and patterns of use vary considerably among individuals. Furthermore, opioid users have identified prescription oxycodone as the drug on which they first became dependent before transitioning to other opioids, including heroin and fentanyl^1-5^. Therefore, this preclinical self-administration paradigm may be capturing the variability within a population where certain individuals have increased oral oxycodone intake. Notably, while we did not assess drug-dependent phenotypes in our present study, operant oral oxycodone self-administration procedures utilizing similar doses and length of self-administration have found drug-dependent phenotypes following oral oxycodone self-administration, indicating oral oxycodone self-administration can result in drug dependence^6^. Future studies investigating whether these profiles are associated with later illicit opioid self-administration are necessary to determine how these preclinical measures relate to patterns reported in humans. Together, these findings support further investigation of the behavioral and neurobiological mechanisms contributing to individual variability in oral oxycodone intake.

Identifying behavioral characteristics associated with subsequent oxycodone intake may improve our understanding of individual vulnerability before opioid exposure. Decision-making performance has previously been associated with oral oxycodone intake, particularly in female rats^23^. Here, we extended those findings by examining behavioral flexibility using a contingency degradation task administered before oral oxycodone exposure. Animals with lower degradation indices, indicative of less flexible behavior, subsequently consumed more oral oxycodone during the last four days of self-administration, whereas animals with higher degradation indices consumed less oxycodone. However, degradation indices were not associated with PC1 scores in either water-or oxycodone-exposed animals. Thus, baseline behavioral flexibility was associated with the amount of oxycodone consumed but not with the broader pattern of self-administration behavior, which loaded onto active responses and reinforcers earned. The relationship accounted for a modest proportion of the variance in oxycodone consumption, and as such, behavioral flexibility as measured by contingency degradation should be considered a potential candidate for determining vulnerability towards oral oxycodone consumption along with other decision-making components or neural factors. Notably, contingency degradation tasks have also been used in human populations^55^, providing a potential framework for future translational studies. The relationship between degradation index and oxycodone consumption was significant in females but not in males, consistent with previous findings that decision-making performance was associated with oral oxycodone intake in female rats^23^. In addition, we have previously shown that estrous-cycle stage can affect contingency degradation performance^41^, which may contribute to variability in females. Therefore, further studies designed to test sex and hormonal status directly will be needed to determine whether baseline behavioral flexibility is differentially related to oxycodone intake in males and females. Lastly, the neural mechanisms shared by contingency degradation performance and oral oxycodone intake have also not been established. Examining whether circuits supporting flexible behavior contribute to variability in subsequent oxycodone consumption may help clarify the mechanisms underlying this association.

Previous studies using operant oral oxycodone self-administration paradigms have reported greater self-administration and addictive-like behaviors in females than in males^6,23^. In the present study, we found no overall sex difference in nose-poke responding or the amount of oral oxycodone consumed, which may reflect differences in the duration of self-administration, oxycodone concentrations, or the use of water rather than a saccharin-fading procedure^23^. While our findings differ from previous operant oral oxycodone self-administration findings^6,23^, they are consistent with short-access intravenous oxycodone studies that reported no effect of sex on oxycodone self-administration^9,56-58^. Hormonal fluctuations across the reproductive cycle have been shown to influence opioid self-administration^9,46,47,59^. Specifically, female rats in proestrus have demonstrated lower opioid self-administration than females in other estrous-cycle stages^9,46,47^. Although nose-poke responding and reinforcers earned did not differ between proestrus and non-proestrus stages during the final four days of self-administration, estrous-cycle stage was associated with the amount of oral oxycodone consumed. These findings suggest that ovarian hormone fluctuations may contribute to variability in oral oxycodone consumption and warrant investigation in future studies.

Substance use disorders affect multiple physiological systems^60^ and orally administered drugs directly contact the gastrointestinal tract. We found that observed richness in fecal samples collected before self-administration was associated with performance on the contingency degradation task. Specifically, greater baseline observed richness was associated with lower degradation indices, indicative of less flexible behavior. This finding is intriguing because relationships between naturally occurring variation in the gut microbiome and baseline behavioral flexibility have not been extensively studied. Previous studies have shown that antibiotic-induced disruption of the microbiome can impair cognitive flexibility in mice^27^ and that dietary manipulations, including high-fat and Western diets, can alter both the gut microbiome and cognitive flexibility^26,61^. The present findings extend previous work by identifying an association between baseline microbiome richness and naturally occurring individual differences in behavioral flexibility. However, this correlational relationship does not establish that microbiome composition influenced behavioral performance. Notably, lower abundance of several of the identified genera was associated with greater behavioral flexibility. Behavioral flexibility was, in turn, negatively associated with oxycodone consumption in the present study, although this relationship was modest, and stronger associations between impaired flexibility and greater drug-taking have been reported elsewhere. Together, these findings raise the possibility that specific microbiome features may track an individual-difference phenotype relevant to vulnerability to elevated drug consumption. However, the present correlational analyses cannot establish whether these microbial differences contribute to behavioral vulnerability, reflect prior behavioral or physiological differences among animals, or emerge because of oxycodone exposure.

Consistent with previous studies examining the effects of opioids on the gut microbiome^24,28,29,39,62^, oral oxycodone exposure was associated with changes in microbiome composition. Intravenous oxycodone studies have shown decreases in beta diversity in fecal samples following self-administration^24^. Our findings have demonstrated similar changes to the gut microbiome in an oral self-administration paradigm where oral oxycodone-exposed animals exhibited lower observed richness following self-administration. Genus-level analyses identified taxa that were nominally associated with behavioral flexibility and taxa that changed following oxycodone exposure. Exploratory genus-level associations identified several low-abundance taxa that were both sensitive to oxycodone exposure and correlated with behavioral flexibility. Because most of these taxa were present at very low relative abundance and have not been directly implicated in behavioral flexibility, these findings should be interpreted as hypothesis-generating rather than evidence of a mechanistic microbiome–behavior relationship. Although some identified genera include host-associated anaerobes, several are primarily characterized in environmental niches, and little is currently known about their potential contributions to behavioral flexibility or opioid-related outcomes. Accordingly, the present findings identify candidate associations rather than genus-specific mechanisms. These taxa should therefore be considered preliminary candidates for future longitudinal studies examining microbiome composition before and after drug exposure.

Together, these findings demonstrate substantial individual variability in oral oxycodone self-administration and identify behavioral and microbiome measures associated with that variability. PCA captured a pattern of self-administration behavior that differed between oxycodone-and water-exposed animals, while hierarchical clustering identified three profiles among oxycodone-exposed animals based on PC1 scores and oxycodone consumption. Baseline behavioral flexibility was associated with subsequent oxycodone consumption in a sex-dependent manner (i.e., driven by female animals), although it was not associated with PC1 and accounted for only a portion of the variability in intake. These behavioral relationships were accompanied by associations between microbiome measures, behavioral flexibility, and oxycodone exposure. Although the present correlational findings cannot establish prediction, causality, or a bidirectional relationship between the gut microbiome and behavior, they identify candidate behavioral and microbial features for future investigation. Future longitudinal and mechanistic studies will be necessary to determine whether these measures contribute to vulnerability to opioid misuse, reflect the consequences of oxycodone exposure, or arise from shared underlying factors.

## ACKNOWLEDGEMENTS

The authors thank Samantha Bozarth for helping run animals during the contingency degradation portion of these experiments, and Dr. Jacqueline-Marie Ferland for help with RNAseq analysis.

## AUTHOR CONTRIBUTION

CC, MN and EW designed the experiments, CC and AO ran the experiments, CC, MN and EW analyzed the data. CC wrote the manuscript, and EW, AO and MN edited the manuscript.

## DATA AVAILABILITY STATEMENT

The datasets generated and analyzed during the current study are available in the GEO repository, GEO accession GSE326997.

