## Supplemental Material for "Behavioral flexibility, gut microbiome composition, and individual variability in oral oxycodone self-administration"

**SUPPLEMENTAL RESULTS.**

**No effect of a higher oral oxycodone dose during early self-administration on oral oxycodone intake during later self-administration.**

Importantly, from days 7-10, a subset of oxycodone-exposed animals (n=12) were maintained at a dose of 1 mg/ml to determine if a higher dose during early self-administration influenced later self-administration behavior. All other oxycodone-exposed animals (n=24) were maintained at a dose of 0.3 mg/ml for days 7-10. As expected, we found that intake between the two groups differed between days 7-10 due to the increase in dose, **but there were no group differences in oxycodone intake prior to self-administration days 1-6 with statistical information provided in Table S5. Importantly, the increase in dose on days 7-10 did not affect oral oxycodone intake during later self-administration (SAD11-21: F(1, 34)=0.001, p=0.9711; Table S1). While we observed a self-administration x dose interaction (F(10,340)=2.063, p=0.02), there were no differences between the two doses on any of the individual days (11-21) between the two groups with posthoc Bonferroni comparisons.**

**No major sex differences during each phase of self-administration.**

We found no major sex differences in active hole and inactive hole responses in oxycodone-exposed and water controls in every phase of self-administration (Fig. S1; SAD1-4, SAD5-13, SAD14-21) **with the statistical information included for active hole and inactive hole responding during self-administration in Table S2 and reinforcers in Table S3.**

**Rats self-administer oral oxycodone with no differences in responding during FR1 schedule of reinforcement compared to water control animals.**

Self-administration was comprised of three phases: 1) self-administration days 1-4 where animals were water regulated to increase motivation to consume liquid reinforcers under a fixed ratio (FR) 1 schedule of reinforcement^33^, 2) self-administration days 5-13 where animals were no longer water regulated, on an FR1 schedule of reinforcement, and had a decrease in dose on self-administration day 10, and 3) self-administration days 14-21 where animals were moved to an FR3 schedule of reinforcement.

**We found that animals poked more into the active hole than the inactive hole during the first two phases of self-administration (SAD1-4: F(1, 57)=380.6, p<0.001; SAD5-13: F(1, 57)=243.3, p<0.001), but there was no difference in these measures between oxycodone-exposed and water control animals (Fig. 2A; SAD1-4: F(1, 57)=1.79, p=0.19; SAD5-13: F(1, 57)=0.046, p=0.83). Additionally, we found no effect of drug exposure on the number of reinforcers earned during self-administration days 1-4 (Fig. 2B; SAD1-4: F(1, 57) = 3.59, p=0.063; SAD5-13: F(1, 57)=0.369, p=0.546). Complete statistical information is provided in Table S4 for active and inactive hole responding and Table S5 for reinforcers earned.**

**All groups have increased responding during the first 30 minutes of the self-administration session and oral oxycodone exposed animals have increased responding overall compared to controls during the last four days of self-administration.**

Time bin analyses (30-minute intervals) on active and inactive hole responding analyses during the last four days of self-administration demonstrated increased responding in oxycodone-exposed animals compared to water animals over the two-hour sessions across the last four days of self-administration (F(1, 234)=580.4, p<0.0001; Fig. S2A; **Table S6**). Oxycodone-exposed animals had increased active hole nose poke responding during the first 30-minute time bin compared to all other time bins (p<0.0001) as did water control animals (30 vs 60, p=0.009, 30 vs 90, p=0.021, 30 vs 120, p=0.002). Time bin analyses conducted on the number of reinforcers earned during the last four days of self-administration demonstrated that oxycodone animals earned more reinforcers than water controls (F(1, 242)=15, p=0.0001; Fig. S2B; **Table S7**). Oxycodone exposed animals earned more reinforcers during the first 30-minute time bin compared to all other time bins (p<0.001) while water control animals earned more reinforcers during the first 30-minute time bin compared to the third (<90, p=0.02) and fourth bins (<120, p=0.01).

**Estrous cycle influences the amount consumed during the last four days of self-administration.**

We did not find a significant main effect of estrous cycle or drug exposure on active and inactive hole nose pokes during self-administration days 18-21, however we did find a significant effect of estrous cycle on reinforcers earned (F(1, 21)=4.747, p=0.0409; Fig. S3). **Statistical analyses on the effects of the estrous cycle on active and inactive hole responses are provided in Table S8 and analyses on reinforcers earned are provided in Table S9.** Importantly when we investigated if there was a difference in the amount of oral oxycodone consumed (n=12) we found that females in the proestrus stage of the estrous cycle have decreased intake compared to when they are in non-proestrus stages of their cycle (Fig. S3C; paired t-test; t(11)=2.423, p=0.0338).

**TABLES.**

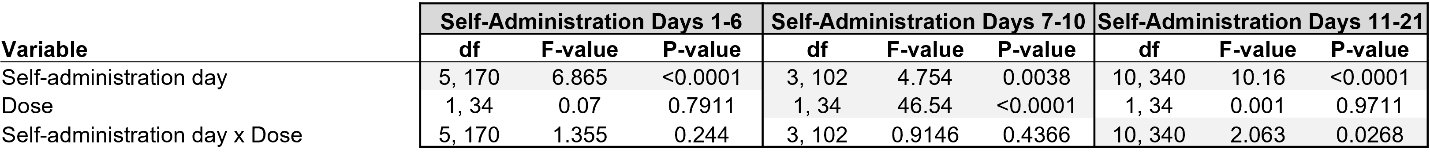

**Table S1. Statistical analysis of amount of oral oxycodone consumed (intake) during self-administration between animals receiving a high dose or a lower dose during self-administration days 7-10.** Two-way mixed ANOVAs were conducted on the amount of oral oxycodone consumed prior to an increase in dose on a subset of animals (n=12; SAD1-6), during the increase in dose (SAD7-10) and later self-administration (SAD11-21). p < 0.05. There were no differences between the two intakes on any of the individual days (11-21) between the two groups with posthoc Bonferroni comparisons. Statistical significances are highlighted in gray.

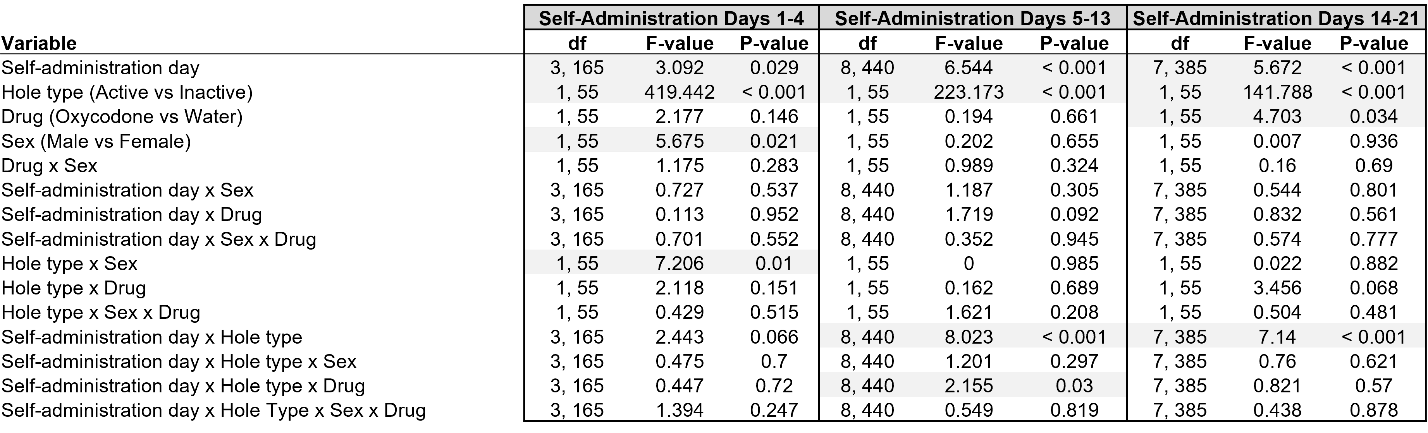
**Table S2. Statistical analysis of active hole and inactive hole responses during each portion of self-administration with sex as a variable.** Four-way mixed ANOVAs were conducted on active and inactive hole responses between the sexes for each phase of self-administration. p < 0.05. Significant effects are highlighted in gray.

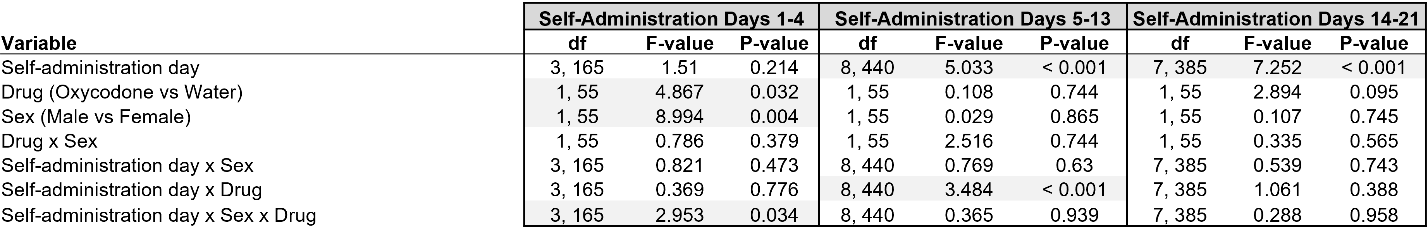

**Table S3. Statistical analysis of reinforcers earned during each portion of self-administration with sex as a variable.** Three-way mixed ANOVAs were conducted on reinforcers earned between the sexes for each phase of self-administration. p < 0.05. Significant effects are highlighted in gray.
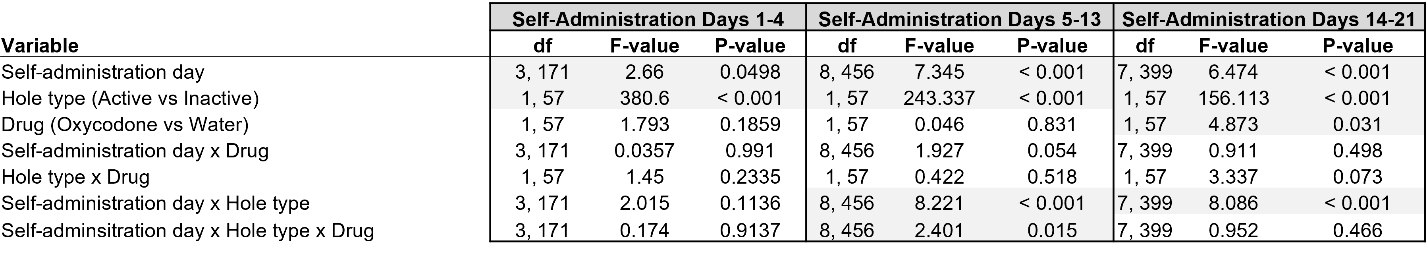

**Table S4. Statistical analysis of active hole and inactive hole responses during each phase of self-administration.** Three-way mixed ANOVAs were conducted on active and inactive hole responses during each phase of self-administration. p < 0.05. Statistical significance are highlighted in gray.

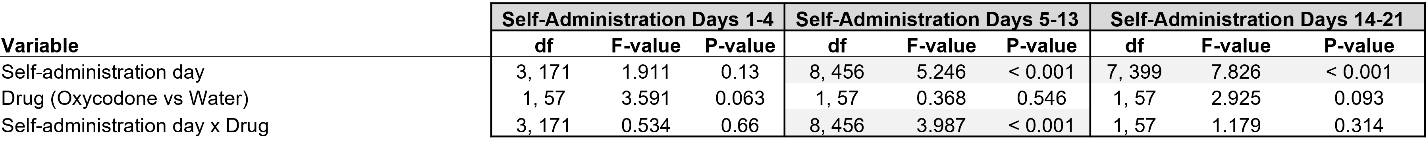

**Table S5. Statistical analysis of reinforcers earned during each phase of self-administration.** Two-way mixed ANOVAs were conducted on the number of reinforcers earned during each phase of self-administration. p < 0.05. Statistical significance are highlighted in gray.

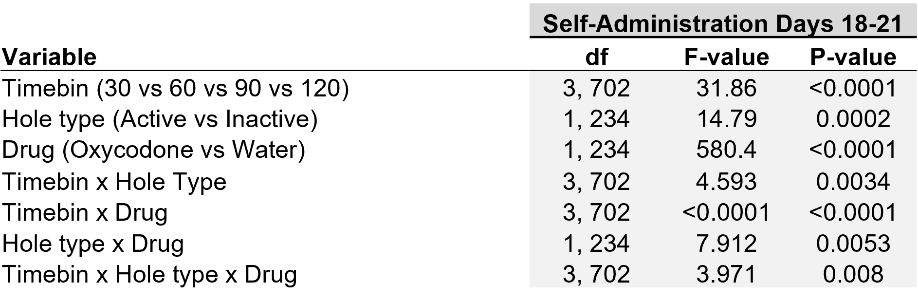

**Table S6. Statistical analysis of active hole and inactive hole responding across 30-minute time bins during the last four days of self-administration (SAD18-21).** Three-way mixed ANOVAs were conducted on active and inactive hole responses from SAD18-21 in oral oxycodone and water control animals during the last four days of self-administration. p < 0.05. Statistical significances are highlighted in gray.

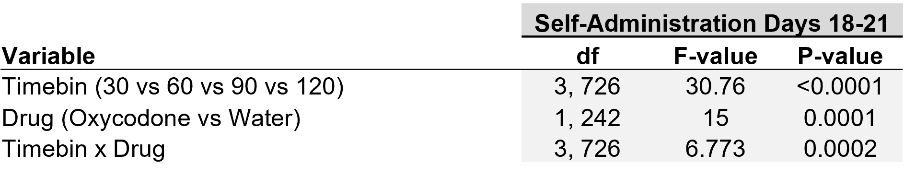

**Table S7. Statistical analysis of reinforcers earned across 30-minute time bins during the last four days of self-administration (SAD18-21).** Two-way mixed ANOVAs were conducted on reinforcers earned from SAD18-21 in oral oxycodone and water control animals during the last four days of self-administration. p < 0.05. Statistical significances are highlighted in gray.

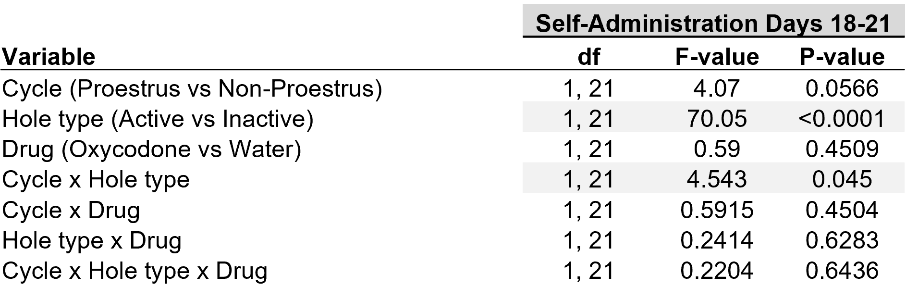

**Table S8. Statistical analysis of the effects of the estrous cycle on active hole and inactive hole responses during the last four days of self-administration (SAD18-21).** Three-way mixed ANOVAs were conducted on active and inactive hole responses from SAD18-21 in oral oxycodone and water control females during the last four days of self-administration. p < 0.05. Statistical significances are highlighted in gray.

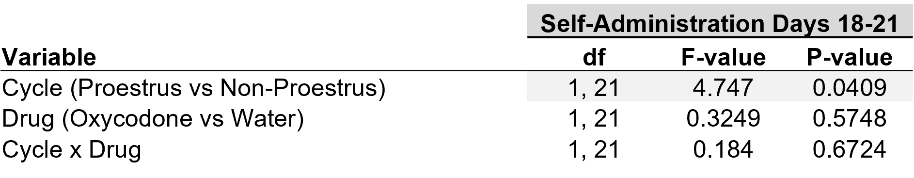

**Table S9. Statistical analysis of estrous cycle effects on the number of reinforcers earned during the last four days of self-administration (SAD18-21).** Two-way mixed ANOVAs were conducted on reinforcers earned from SAD18-21 in oral oxycodone and water control females during the last four days of self-administration. p < 0.05. Statistical significances are highlighted in gray.

**FIGURES.**

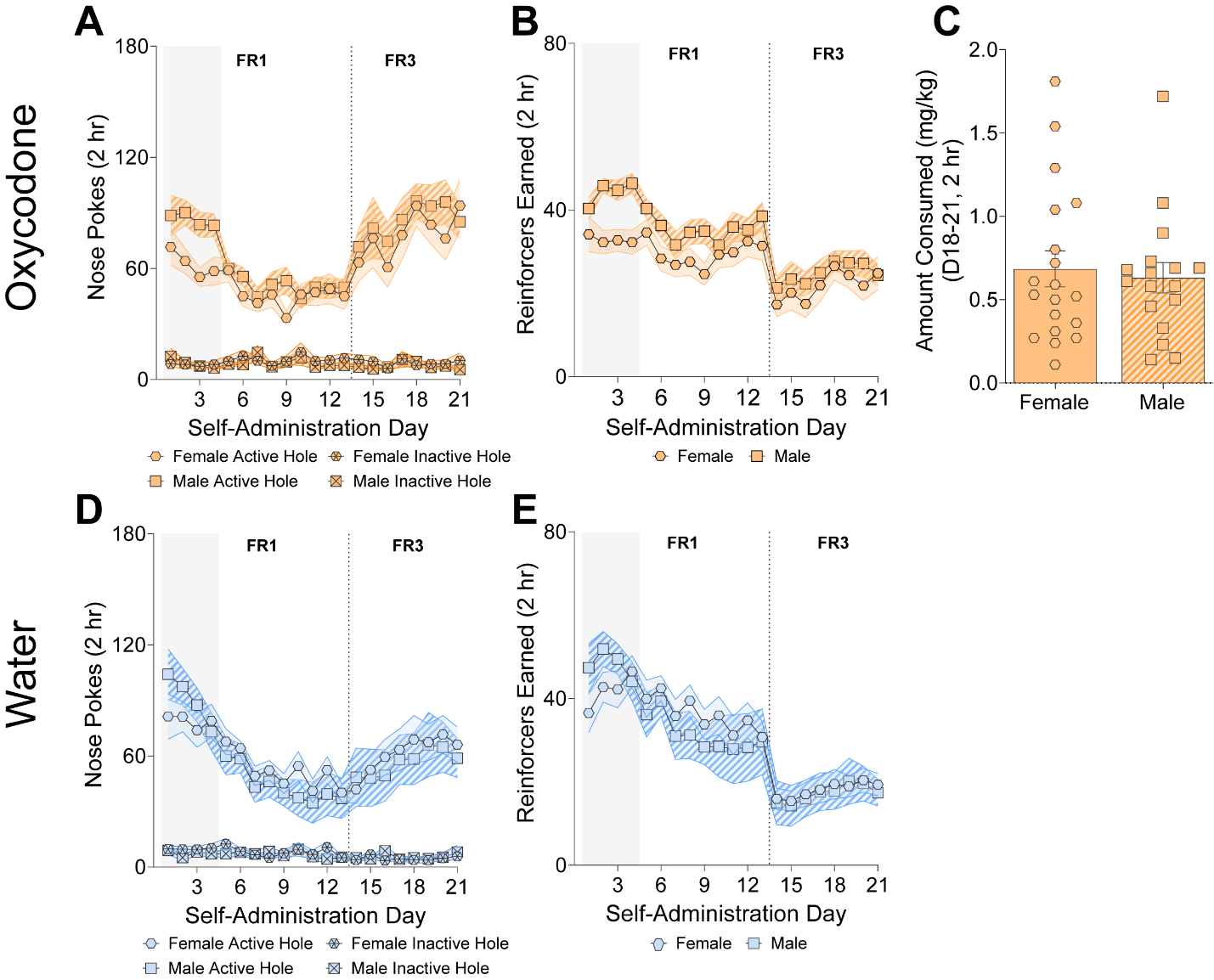

**Figure S1. No major sex differences in nose poke responding and reinforcers earned in oral oxycodone and water control animals or in oral oxycodone consumed in oxycodone exposed animals. A,D.** Males responded more during the first four days of self-administration compared to females (four-way ANOVA, SAD1-4: F(1, 55) = 5.675, p=0.02) but there were no differences in responding between the sexes in the second and third phases of self-administration (SAD5-13: F(1, 55)=0.20, p=0.66; SAD14-21: F(1, 55)=0.007, p=0.94). **B,E.** There was a significant difference in the number of reinforcers earned during the first four days of self-administration between female and male animals (three-way ANOVA, F(1, 55)=8.994, p = 0.004). **C.** There was no difference in the amount consumed during the last four days of self-administration between males and females (unpaired t-test; t(34)=0.37, p=0.71). Data are expressed as mean ± SEM. n=15 water female, n=8 water male; n=19 oxycodone female, n=17 oxycodone male.

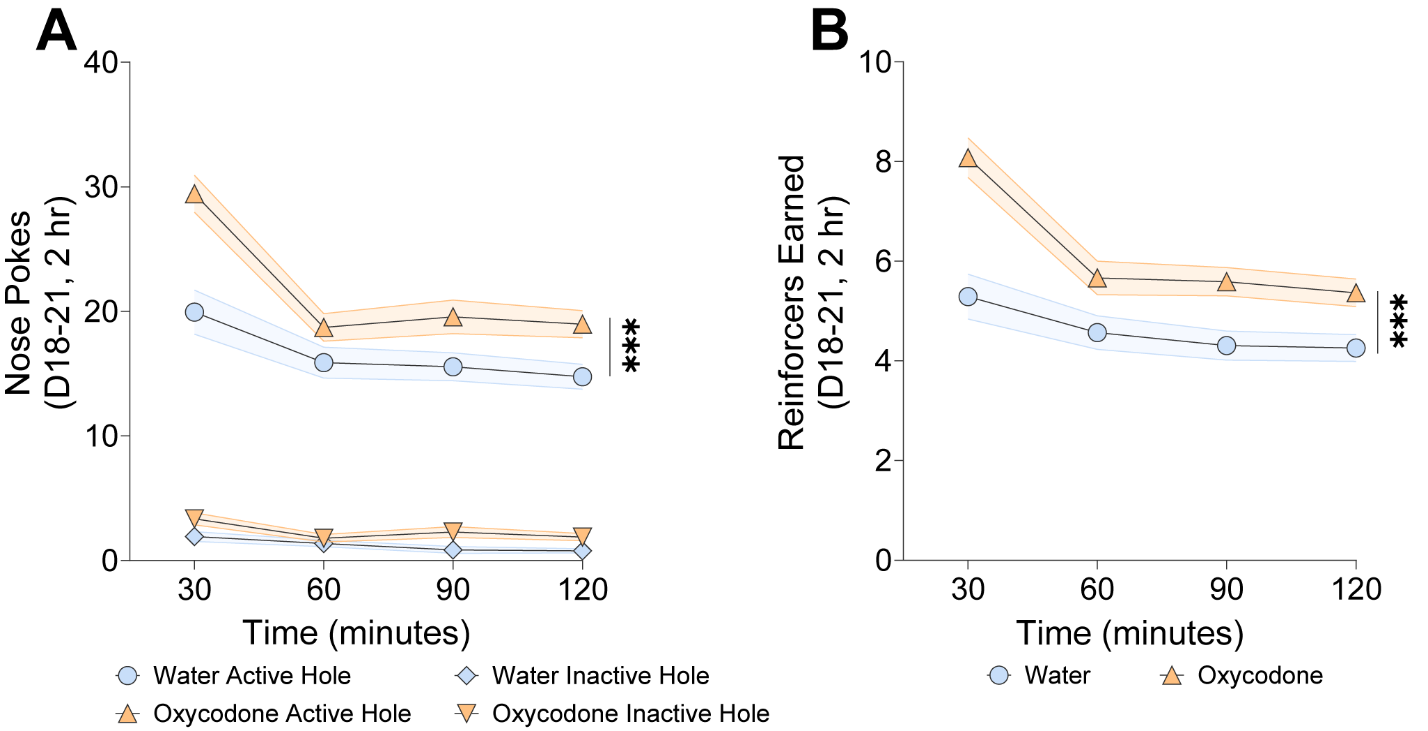

**Figure S2. Oral oxycodone and water animals increase active hole nose pokes and earn more reinforcers during the first 30 minutes of a session during the last four days of self-administration. A.** Oxycodone exposed animals have increased active hole responding during each 30-minute interval compared to water controls during the last four days of self-administration (three-way ANOVA; oxycodone vs water, F(1, 234)=580.4, p<0.0001; active vs inactive, F(1, 234) = 14.79, p=0.0002). Both groups have increased responding during the first 30 minutes of the session (oxycodone: 0-30 min vs all other time bins, p < 0.0001; water: vs <60 min, p<0.01, vs <90 min, p=0.021, <120 min, p=0.002). **B.** Oxycodone exposed animals earn more reinforcers during the first 30-minute interval compared to all other time bins (p<0.001) while water control animals earned more reinforcers during the first 30-minute time bin compared to the third (<90, p=0.02) and fourth bins (<120, p=0.01). Data are expressed as mean ± SEM. n=23 water, 36 oxycodone.

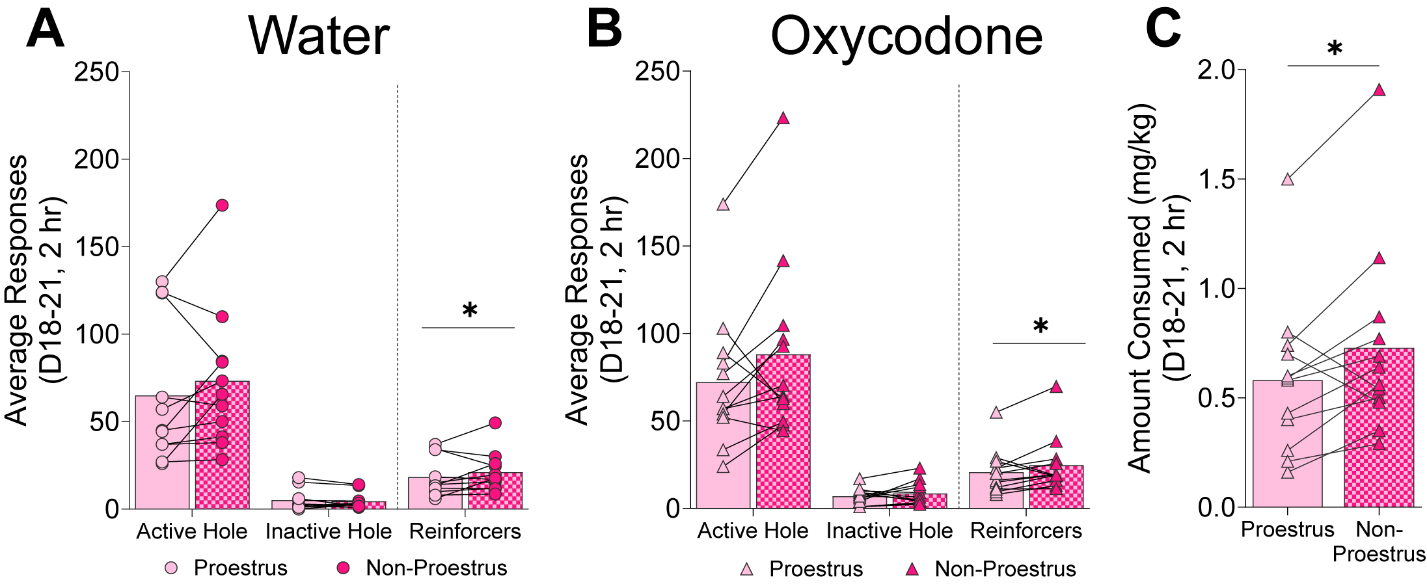

**Figure S3. Estrous cycle influences the amount of reinforcers earned in water and oral oxycodone-exposed animals and the amount oral oxycodone consumed during the last four days of self-administration. A,B.** Active and inactive nose pokes responses and reinforcers earned during the last four days of self-administration (SAD18-21) in water control **(A)** and oral oxycodone **(B)** females in proestrus and non-proestrus stages of the estrous cycle. There was no effect of estrous cycle (three-way ANOVA; F(1, 21)=4.07, p=0.0566) or drug exposure (F(1, 21)=0.59, p=0.4509) on active and inactive hole responses. There was an effect of estrous cycle on the number of reinforcers earned (F(1, 21) = 4.747, *p = 0.0409). **C.** Females in the proestrus stage of the estrous cycle consumed less oral oxycodone compared to when the same female was in non-proestrus stages during SAD18-21 (paired t-test; t(11) = 2.423, p = 0.0338). Data are expressed as mean ± SEM. n=11 water, n=12 oxycodone.
